# Differentiating Hepatic and Renal Toxicity Reveals CYP-Independent Mechanisms of Acetaminophen-Induced Acute Kidney Injury

**DOI:** 10.64898/2026.06.15.732380

**Authors:** Yasaman Etemadi, Timothy A. Fields, Anup Ramachandran, Hartmut Jaeschke

## Abstract

Acetaminophen (APAP) overdose is the leading cause of acute liver failure (ALF), with acute kidney injury (AKI) contributing substantially to morbidity and mortality in those patients. To determine whether APAP-induced AKI depends on hepatic CYP2E1-mediated bioactivation, we used CYP2E1^flox/flox^ mice treated with AAV8-TBG-Cre to selectively delete hepatic CYP2E1 while preserving renal metabolism. Male and female mice received APAP (600 mg/kg) and were evaluated up to 48 hours for liver and kidney injury. Liver-specific CYP2E1 deletion reduced APAP hepatotoxicity, confirming the absence of hepatic NAPQI formation. Despite this protection, both male and female mice treated with AAV8-TBG-Cre and APAP developed progressive renal injury, with marked increases in blood urea nitrogen (BUN) and creatinine, tubular vacuolation, and strong induction of KIM-1 and osteopontin, along with apoptotic cell death at 48 hours. Notably, female mice, lacking renal CYP2E1 and displaying no detectable renal protein adducts, still progressed to AKI, demonstrating that kidney injury can occur through CYP-independent mechanisms. Given that APAP-induced AKI is a delayed injury, we further considered p-aminophenol (PAP), a deacetylation product of APAP, as a potential CYP-independent contributor. These findings support the concept that non-CYP pathways, including PAP formation, may contribute to kidney injury during the later phase of toxicity, although this pathway likely represents only one component of a multifactorial injury process. Together, these results demonstrate that APAP-induced AKI is a kidney-intrinsic process that can develop independently of both hepatic and renal CYP2E1 activity, emphasizing the need for kidney-specific therapeutic strategies for preventing APAP-induced renal injury.

## 1. Introduction

Acetaminophen (APAP) is one of the most widely used over-the-counter analgesics and antipyretics, yet overdose remains the leading cause of acute liver failure (ALF) in the United States and other Western countries (Lee, 2004). APAP-related ALF accounts for nearly half of all liver failure cases in the US and continues to drive substantial emergency department visits, hospitalizations, and mortality (Hodgman and Garrard, 2012; Larson et al., 2005; Ostapowicz et al., 2002). Although the mechanisms of APAP-induced hepatotoxicity have been extensively characterized (Jaeschke and Ramachandran, 2024a; Ramachandran and Jaeschke, 2019), progression to multi-organ injury remains a major clinical challenge (Chen et al., 2015; Mazer and Perrone, 2008; Stollings et al., 2016; Tujios et al., 2015). Acute kidney injury (AKI) is particularly important in this context: clinical studies report that 30–50% of ALF patients develop AKI, and elevations of renal biomarkers such as BUN and creatinine correlate with more severe renal dysfunction and poorer survival (Antoine et al., 2015; Hadem et al., 2019; Tujios et al., 2015). Patients who develop AKI also exhibit significantly lower one-year survival rates (Tujios et al., 2015). Consistent with this, our prior clinical analysis showed that APAP overdose patients who developed renal failure consistently had more severe hepatic injury than those without renal involvement (Akakpo et al., 2020).

During APAP overdose, a large fraction of the drug is bioactivated by cytochrome P450 2E1 (CYP2E1) to the reactive metabolite N-acetyl-p-benzoquinone imine (NAPQI) (Jaeschke and Ramachandran, 2024b; McGill and Jaeschke, 2013). Excessive NAPQI formation rapidly depletes hepatic glutathione, disrupts mitochondrial function, and ultimately drives hepatocellular necrosis (Gujral et al., 2002; Jaeschke et al., 2019). In contrast to the liver, the early mechanisms of APAP-induced kidney injury are fundamentally different. Although CYP2E1 in proximal tubular cells contributes to local NAPQI formation (Akakpo et al., 2020; Hart et al., 1994, 1995), renal injury does not primarily depend on mitochondrial dysfunction (Akakpo et al., 2023). Instead, accumulating evidence—supported by our recent studies—indicates that endoplasmic reticulum (ER) stress and caspase-mediated apoptosis are central drivers of APAP nephrotoxicity (Akakpo et al., 2023; Lorz et al., 2004). Consistent with this, the caspase inhibitor Ac-DEVD-CHO selectively attenuates kidney injury without altering liver damage, demonstrating distinct organ-specific toxicity pathways (Akakpo et al., 2023). Notably, although NAC effectively mitigates hepatic injury by replenishing GSH, it provides limited renal protection, revealing a major therapeutic gap (Davenport and Finn, 1988; Hengy et al., 2009; Jones and Vale, 1993; Slitt et al., 2004).

Despite the clinical importance of APAP-induced AKI and the distinct injury mechanisms between liver and kidney, the mechanistic relationship between hepatic metabolism and renal injury remains unresolved. Previous studies have been unable to separate hepatic from renal contributions because they relied on whole-body knockouts, systemic inhibitors, or otherwise non-selective approaches (Emeigh Hart et al., 1996; Lee et al., 1996; Slitt et al., 2005). Although emerging evidence suggests potential liver–kidney crosstalk, including microRNA-mediated regulation of renal CYP2E1 (Matthews et al., 2020), several clinical reports indicate that APAP-induced nephrotoxicity can occur independently of overt hepatotoxicity (Campbell and Baylis, 1992; Hadem et al., 2019; Leithead et al., 2009). Together, these limitations highlight a critical gap: whether AKI can develop in the absence of hepatic NAPQI formation. Addressing this question requires a model in which hepatic or renal oxidative APAP metabolism is selectively eliminated.

An additional layer of complexity to interpreting APAP-induced nephrotoxicity are sex differences in CYP2E1 expression. Although males generally exhibit higher CYP2E1 expression in the kidney and activity than females, this difference does not translate into a consistent sex-specific susceptibility to APAP-induced AKI in clinical settings (Hoivik et al., 1995; Hu et al., 1993). This discrepancy raises an important unresolved question: do observed sex differences arise primarily from hepatic metabolism, renal bioactivation, or a combination of both? Distinguishing these factors requires an experimental model in which hepatic CYP2E1 is selectively eliminated while renal CYP2E1 expression is preserved, enabling direct assessment of kidney-intrinsic mechanisms of APAP bioactivation and injury. Adeno-associated virus serotype 8 (AAV8)-TBG-Cre, which pairs the inherent hepatotropism of AAV8 with the liver-restricted thyroid hormone–binding globulin (TBG) promoter, enables highly efficient and selective Cre-mediated recombination in hepatocytes with the floxed allele (Kiourtis et al., 2021). This technique enables us to test whether APAP-induced AKI persists when CYP2E1-mediated APAP metabolism and NAPQI formation are eliminated in the liver. Thus, resolving whether APAP-induced AKI can occur independently of liver injury is essential for defining kidney-specific injury pathways and guiding the development of therapeutic strategies that extend beyond NAC to improve renal outcomes after APAP overdose.

## 2. Materials and methods

### 2.1. Animals

Male and female C57BL/6J mice (8–10 weeks old) were purchased from Jackson Laboratories (Bar Harbor, ME, USA) and maintained under standard housing conditions with ad libitum access to food and water. Heterozygous Cyp2e1 floxed mice on the C57BL/6J background (Cyp2e1^flox/+^) were obtained from Cyagen (Santa Clara, CA) and intercrossed to generate homozygous Cyp2e1^flox/flox^ mice. Genotypes were confirmed by PCR analysis of genomic DNA isolated from tail biopsies based on the genotyping protocol from Cyagen. Homozygous floxed mice were used for all subsequent experiments. Animals were kept in a controlled environment with regulated temperature and a 12-hour light–dark cycle. All procedures were conducted in accordance with institutional and federal guidelines and approved by the University of Kansas Medical Center Institutional Animal Care and Use Committee.

### 2.2. Experimental design

All reagents were obtained from Sigma-Aldrich (St. Louis, MO) unless otherwise specified. Mice were fasted for 16 hours overnight before treatment. APAP was dissolved in warmed saline and administered intraperitoneally (i.p.) at a dose of 600 mg/kg. For AAV8 experiments, mice received an intravenous (i.v.) injection of AAV8-TBG-Cre or AAV8-null control (1 × 10¹¹ genome copies in saline) two weeks before APAP administration. Mice were euthanized 24 or 48 hours after APAP treatment under isoflurane anesthesia. In a separate experiment, AAV8-Cre–treated male mice were pretreated with 125 mg/kg Tri-O-tolyl phosphate (TOTP) dissolved in corn oil 1 hour before APAP (600 mg/kg) and sacrificed 48 hours after treatment. Blood was collected from the inferior vena cava using heparinized syringes and centrifuged at 18,000 × g for 2 minutes to isolate plasma. Livers and kidneys were excised, rinsed in saline, and processed. Tissues were fixed overnight in 10% phosphate-buffered formalin for histological analysis, whereas the remaining samples were snap-frozen in liquid nitrogen for biochemical and molecular assays.

### 2.3. Biochemical assays

Plasma alanine aminotransferase (ALT) levels were measured using a commercial colorimetric assay kit (MedTest, Canton, MI, USA) according to the manufacturer’s instructions. Kidney function was assessed by quantifying BUN and creatinine using the QuantiChrom™ Urea Assay and QuantiChrom™ Creatinine Assay kits (BioAssay Systems, Hayward, CA), following the standardized protocols provided by the manufacturer.

### 2.4. Histology and Immunohistochemistry

For histological analysis, liver and kidney tissues were fixed in neutral buffered formalin solution before undergoing paraffin embedding. Embedded tissue blocks were sectioned at a 5-micron thickness using a microtome. To assess morphological features and quantify necrotic regions, mounted tissue sections were stained with hematoxylin and eosin (H&E) or Periodic Acid–Schiff (PAS) staining according to standard protocols. DNA fragmentation in kidney sections was assessed using terminal deoxynucleotidyl transferase-mediated dUTP nick end-labeling (TUNEL) staining. The staining was performed with the In-Situ Cell Death Detection Kit-fluorescein (Roche Diagnostics, Indianapolis, IN; Cat#11684795910). For Kidney Injury Marker1 (KIM1) and osteopontin immunostaining, kidney sections were first blocked for 1 hour at room temperature with 3% BSA in serum and then incubated overnight with rabbit anti-KIM1 (Invitrogen #PA5-46914) or rabbit anti-osteopontin antibodies (Cell Signaling Technology, 66614). To quench endogenous peroxidase activity, sections were treated with 3% hydrogen peroxide, followed by three washes in TBS (10 minutes each). Signal detection was carried out using an avidin-biotin conjugate detection system (Vector Laboratories, PK-4001) followed by 3,3′-diaminobenzidine (DAB) substrate kit (Cell Signaling Technology, 8059). For selected experiments, sections were first immunostained for osteopontin and subsequently counterstained with PAS (Persy et al., 1999) to distinguish expression in proximal versus distal tubules after APAP overdose.

### 2.5. Western Blotting

Western blot analysis was performed to assess CYP2E1 expression. Proteins were separated by SDS–PAGE and transferred onto membranes, which were blocked with 5% BSA and incubated overnight at 4°C with primary antibodies (1:1000): rabbit anti-CYP2E1 (Abcam, ab28146) and rabbit anti–β-actin (Cell Signaling Technology #4970). Membranes were subsequently incubated with HRP-conjugated secondary antibody (goat anti-rabbit, 1:2000; Invitrogen #32460) and developed using chemiluminescent detection reagents (Cytiva, Marlborough, MA). Signals were captured using a LI-COR Odyssey imaging system (LI-COR Biosciences, Lincoln, NE).

### 2.6. RNA isolation and quantitative PCR

Total RNA was isolated from liver and kidney tissues using TRIZOL reagent (Invitrogen) per the manufacturer’s protocol. Two micrograms of purified RNA were reverse transcribed into cDNA using a commercially available kit (Applied Biosystems #4368814). Quantitative PCR was performed using PowerUp SYBR Green Master Mix (ThermoFisher #A25742) with gene-specific primers purchased from Integrated DNA Technologies. The mRNA expression level of CYP2E1 was quantified by the comparative CT method using 18S rRNA as an internal reference gene. Relative fold-changes in gene expression were calculated using the 2−ΔΔCT method.

### 2.7. Quantitation of APAP protein adducts

Protein adducts were analyzed following the methods described previously (Etemadi et al., 2023; McGill et al., 2013). In brief, kidney tissue homogenates were filtered using Bio-Spin 6 columns (Bio-Rad, Hercules, CA) to eliminate low molecular weight metabolites that could interfere with the detection of APAP protein adducts. Tissue proteins were precipitated and then digested with proteases. After additional centrifugation, the isolated APAP-CYS residues were analyzed via high-pressure liquid chromatography (HPLC) equipped with a Coularray electrochemical detector (ESA Biosciences, Chelmsford, MA).

### 2.8. Quantitation of P-Aminophenol (PAP) By Alkaline Phenol Assay

PAP levels were measured as an indicator of CYP-independent metabolism contributing to APAP-induced acute kidney injury. Briefly, 100 mg of liver and kidney tissue were homogenized in 0.5 mL of lysis buffer (25 mM HEPES, 5 mM EDTA, 0.19% CHAPS) containing protease inhibitors. The homogenates were centrifuged at 10,000 × g for 20 min, and the supernatants were collected for analysis of APAP deacetylation activity. Protein concentration in the lysates was determined using a BCA assay (Thermo Fisher Scientific, Waltham, MA). For PAP measurement, 100 μL of tissue lysate was mixed with 50 μL of ice-cold trichloroacetic acid (TCA) to precipitate proteins. Samples were centrifuged at 10,000 × g for 10 min, and the resulting supernatants were collected. An equal volume of 5% phenol in 2.5% NaOH and 2.5% Na_2_CO_3_ solutions was then added, and samples were incubated at room temperature for 20 min. Absorbance was measured at 630 nm to quantify PAP levels. PAP concentrations were normalized to total protein content for each sample (Emeigh Hart et al., 1991).

### 2.9. Statistical analysis

Quantitative data are presented as mean ± standard error of the mean (SEM). Statistical comparisons between two experimental groups were conducted using an unpaired Student’s T-test. Comparisons between multiple groups were performed using one-way ANOVA followed by Student-Newman-Keuls post hoc test for multiple groups. Differences with a p-value less than 0.05 were considered to be statistically significant. All graphical representations and analyses were performed using GraphPad Prism version 8.0.1 (GraphPad Software, San Diego, CA).

## 3. Results

### 3.1. Administration of a severe APAP overdose results in pronounced acute kidney injury

To characterize organ injury following severe APAP overdose, male C57BL/6J mice were administered 600 mg/kg APAP and evaluated 24 hours later. In our previous work, we demonstrated that a moderate APAP dose (300 mg/kg) produces measurable hepatotoxicity but does not cause acute kidney injury at any time points (Etemadi et al., 2026), whereas the severe 600 mg/kg dose induces clear renal dysfunction (Akakpo et al., 2024; Akakpo et al., 2023). Therefore, the 600 mg/kg dose was selected for detailed assessment of kidney-specific injury mechanisms. Consistent with substantial hepatic damage, serum ALT levels were markedly elevated compared to controls (Figure 1A). BUN and creatinine levels were also significantly increased, confirming functional impairment of the kidney at this higher dose (Figure 1B, C). Histological examination of liver tissue demonstrated widespread centrilobular necrosis in the APAP-treated group compared to the control group (Figure 1D).

**Figure 1:**
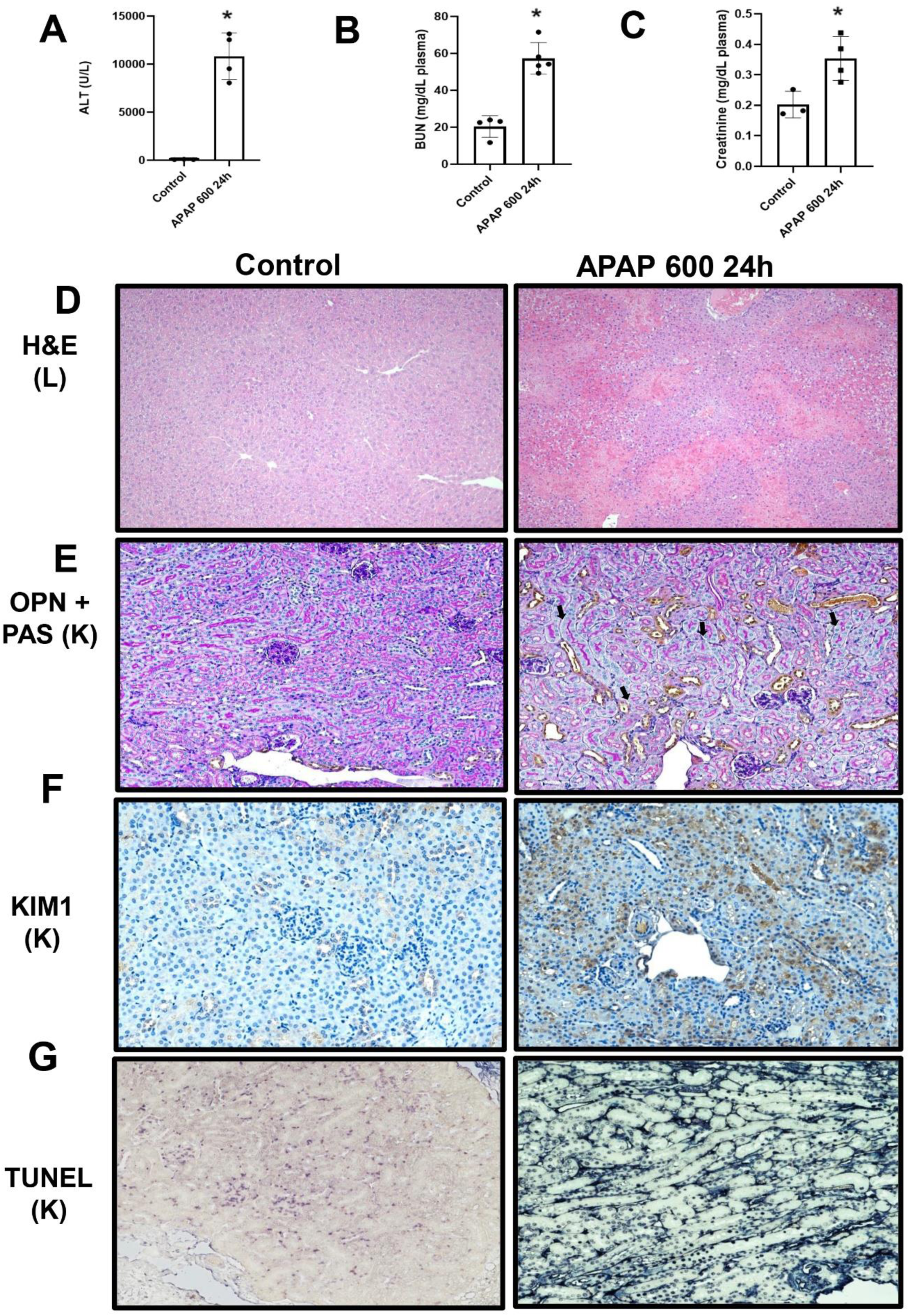
APAP 600 mg/kg induces acute kidney and liver injury at 24 hours. Fasted male C57BL/6J mice received a single intraperitoneal injection of APAP (600 mg/kg) and were sacrificed 24 hours later. (A) Plasma ALT activities, (B) plasma BUN levels, and (C) plasma creatinine levels, (D) Liver H&E staining (10X), (E) Kidney OPN co-staining with PAS counterstain (20X), (F) Kidney KIM-1 immunohistochemistry (20X), (G) Kidney TUNEL staining (20X). Bars represent means ± SEM for n = 4–5 mice per group. *p < 0.05 compared to control.

Histopathological analysis also indicated extensive kidney damage after severe APAP overdose. PAS staining, which highlights the brush border of proximal tubules and distinguishes proximal from distal segments (Gaut and Liapis, 2021; Kellum et al., 2013), showed prominent vacuolar formation in proximal tubular epithelial cells (arrows), indicating acute kidney injury (Figure 1E). To further localize tubular damage, we performed osteopontin immunolabeling. OPN is a secreted glycoprotein expressed in injured renal tubules and serves as a sensitive marker of tubular stress and epithelial damage (Cen et al., 2017; Khamissi et al., 2022; Xie et al., 2001). Combining OPN immunolabeling with PAS staining enabled clear identification of segment-specific injury (Figure 1E). In control cortex, OPN staining was not visible in any cell type. In APAP-treated mice, in the cortex, OPN induction was confined to distal tubules, being absent in proximal tubules with PAS-positive brush borders (Figure 1E). KIM-1 immunostaining provided complementary evidence of injury in proximal tubules; KIM-1 is a highly sensitive and specific biomarker of proximal tubular epithelial injury (Tanase et al., 2019; Wajda et al., 2020), and APAP-treated kidneys exhibited intense KIM-1 expression throughout injured proximal tubule cells compared to the controls (Figure 1F). Finally, TUNEL staining identified extensive DNA fragmentation in renal tissue following APAP overdose (Figure 1G).

### 3.2. Hepatic CYP2E1 deletion reduces liver injury while the kidney remains susceptible

To determine how hepatic APAP metabolism influences kidney injury—and to prevent fatal liver failure that would otherwise preclude consistent evaluation of mice treated with severe APAP overdose beyond 24 hours—we used AAV8-TBG-Cre (Bhushan et al., 2021; Yanger et al., 2014) to selectively delete CYP2E1 in hepatocytes of CYP2E1^flox/flox^ mice. We hypothesized that this approach would significantly reduce liver-specific APAP bioactivation via CYP2E1 and improve survival while preserving renal CYP2E1 expression, thereby enabling assessment of kidney injury mechanisms independent of hepatic metabolism and facilitating investigation of APAP-induced AKI beyond 24 h. As shown in Figure 2A, AAV8-TBG-Cre treatment resulted in selective hepatic deletion of CYP2E1, with preserved CYP2E1 expression in the kidney. Serum ALT levels were markedly elevated in AAV8-null mice given 600 mg/kg APAP but were dramatically reduced in AAV8-TBG-Cre mice at both 24 and 48 hours (Figure 2B), confirming effective protection from hepatotoxicity. In contrast, BUN and creatinine remained significantly elevated in both groups after APAP administration (Figure 2C, D), demonstrating that renal dysfunction persists despite extensive reduction of hepatic APAP toxicity. Western blotting verified efficient hepatic CYP2E1 deletion in AAV8-TBG-Cre mice after APAP treatment, with no re-expression at 24 hours (Figure 2E). Immunohistochemistry confirmed these findings: hepatic CYP2E1 staining was absent in AAV8-TBG-Cre mice at 24-48h but preserved in AAV8-null controls following APAP overdose (Figure 2F). Together, these results confirm sustained hepatocyte-specific CYP2E1 deletion with AAV8 treatment and show that severe APAP overdose produces kidney injury independent of hepatic bioactivation.

**Figure 2:**
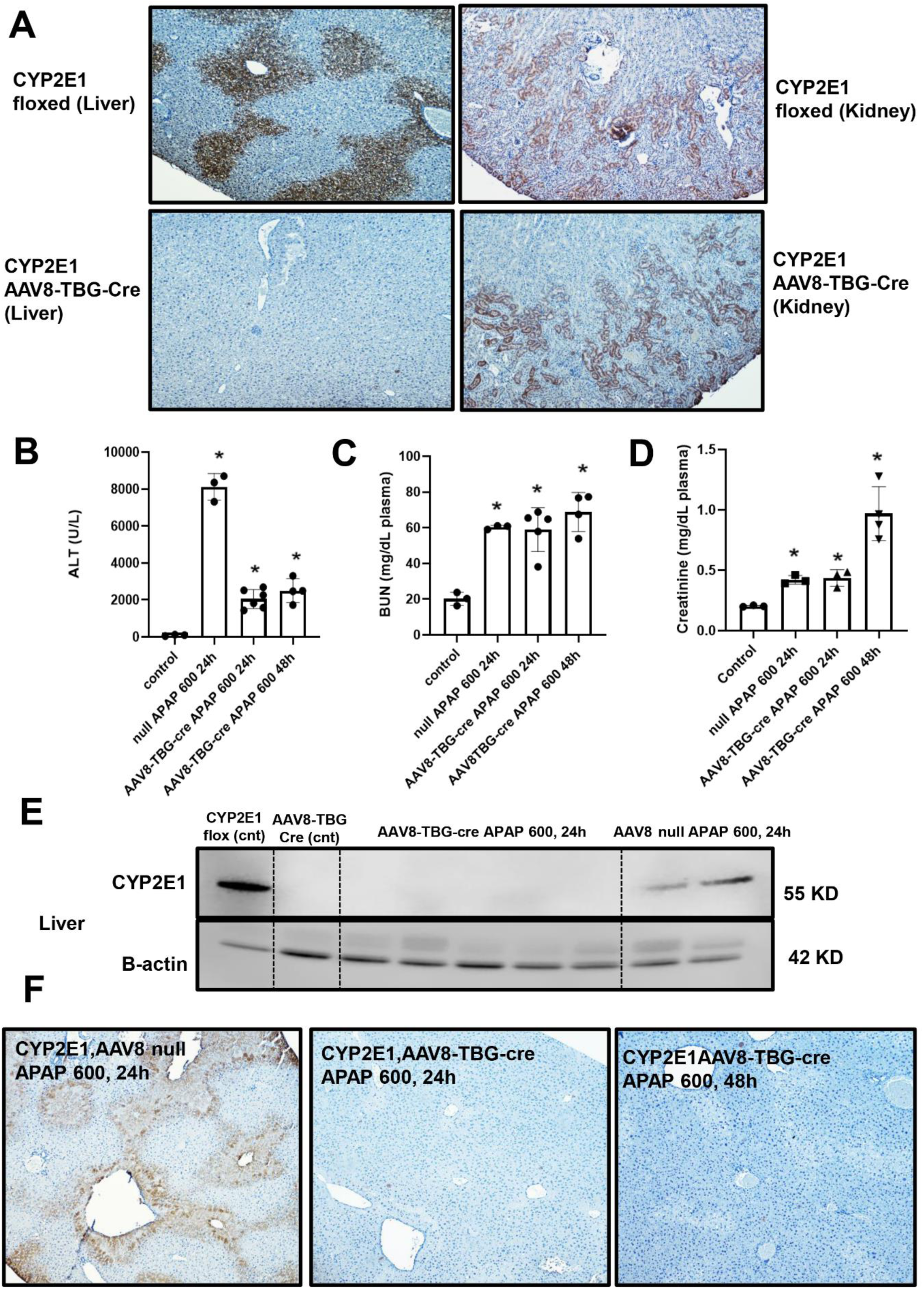
Selective hepatic CYP2E1 deletion uncovers liver-independent mechanisms of APAP nephrotoxicity. CYP2E1-floxed male C57BL/6J mice were administered AAV8-TBG-Cre or AAV8-null vectors two weeks before receiving APAP (600 mg/kg) and were sacrificed 24 or 48 hours later. (A) Liver-specific deletion of CYP2E1 was confirmed by immunohistochemical staining for CYP2E1 in liver and kidney sections, (B) plasma ALT activities, (C) plasma BUN levels, (D) plasma creatinine levels, (E) liver CYP2E1 protein level assessed by Western blot, (F) liver CYP2E1 immunohistochemistry (10X) in AAV8-null or AAV8-TBG-Cre mice following APAP treatment for 24 or 48 hours. Bars represent means ± SEM for n = 4–5 mice per group. *p < 0.05 compared to control.

Histological analysis of liver and kidney sections further supported these findings (Figure 3). Consistent with the biochemical markers of hepatocellular injury shown in Figure 2A, liver sections from AAV8-TBG-Cre treated mice exhibited limited necrosis at either 24- or 48-hours following APAP administration, confirming effective protection from hepatic damage. In contrast, kidney histology revealed substantial renal injury in mice receiving AAV8-null or Cre following 600 mg/kg APAP. At 24 hours, proximal tubular cells displayed prominent cytoplasmic vacuolization (Black arrow), indicative of early tubular injury. Notably, by 48 hours, kidneys from AAV8-TBG-Cre treated mice exhibited marked tubular dilation and abundant intraluminal casts (Yellow arrow) consisting of Tamm–Horsfall protein (uromodulin) (Iorember & Vehaskari, 2014), indicating progression to severe and irreversible tubular injury (Figure 3). These findings were independently confirmed by an expert renal pathologist (TF). In Figure 4, kidney injury was further assessed in AAV8-treated mice following APAP overdose using KIM-1, osteopontin, and TUNEL staining across all groups. At 24 hours, mice receiving AAV8-null and AAV8-TBG-Cre showed comparable levels of KIM-1 and osteopontin expression, as well as similar apoptotic cell death, consistent with early tubular injury. However, by 48 hours, AAV8-TBG-Cre–treated mice exhibited a marked increase in all three markers—KIM-1, osteopontin, and TUNEL-positive cells—indicating substantial worsening of tubular injury and DNA fragmentation following APAP exposure. This delayed but pronounced rise in injury markers aligns with the progression to irreversible renal damage evident in histological analyses.

**Figure 3:**
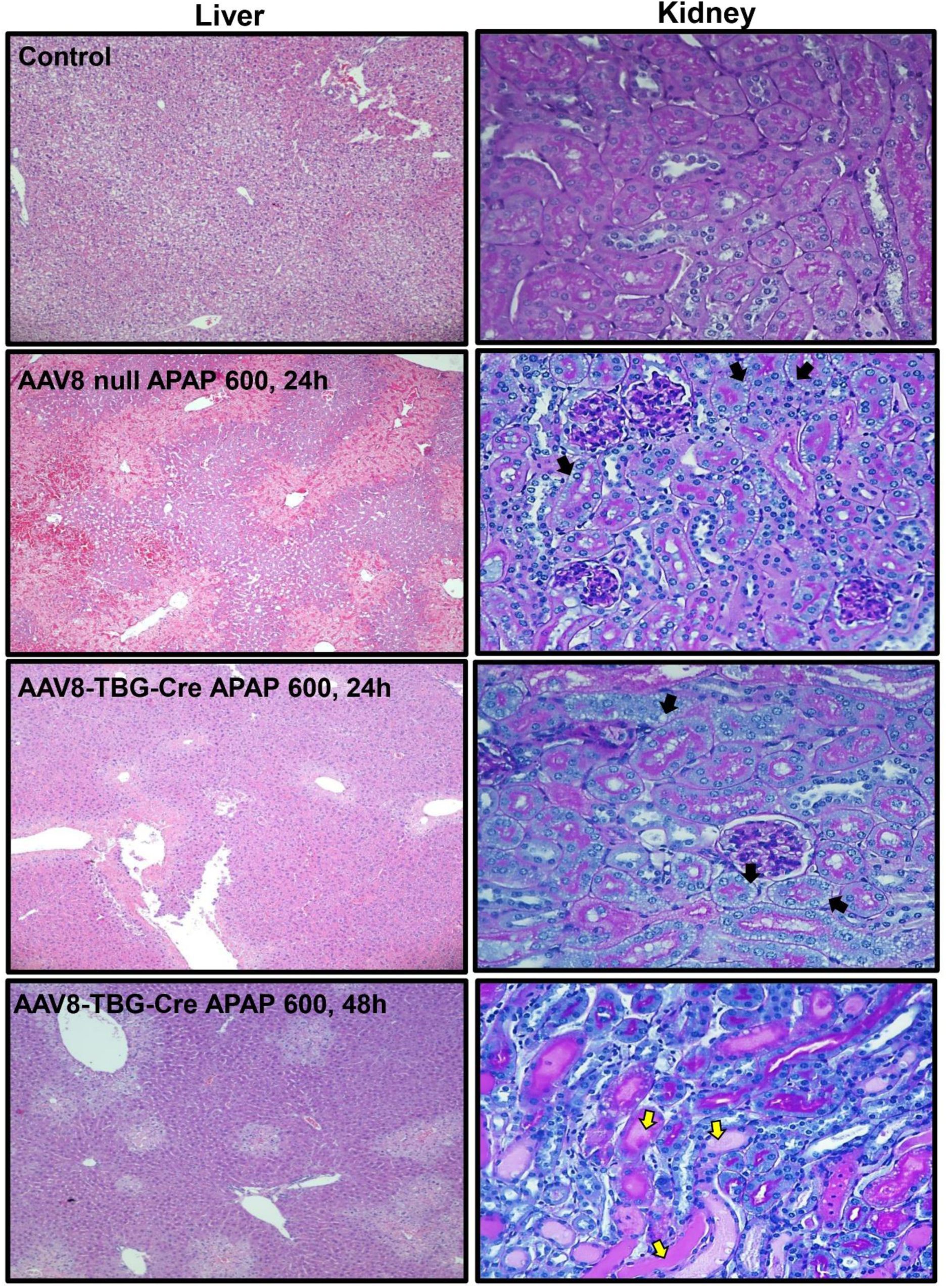
Progressive morphological disruptions in renal tissues following selective hepatic CYP2E1 deletion. Liver (left panels,10X) and kidney (right panels,40X) sections were stained with H&E and PAS to evaluate tissue morphology. Representative liver sections are shown for untreated mice and for mice examined 24 or 48 hours after APAP treatment. Corresponding kidney sections show cortical morphology under the same treatment conditions, with black arrows marking areas of vacuolar change in tubules and yellow arrows indicating tubular cast formation. Images are representative of n = 4–5 mice per group.

**Figure 4:**
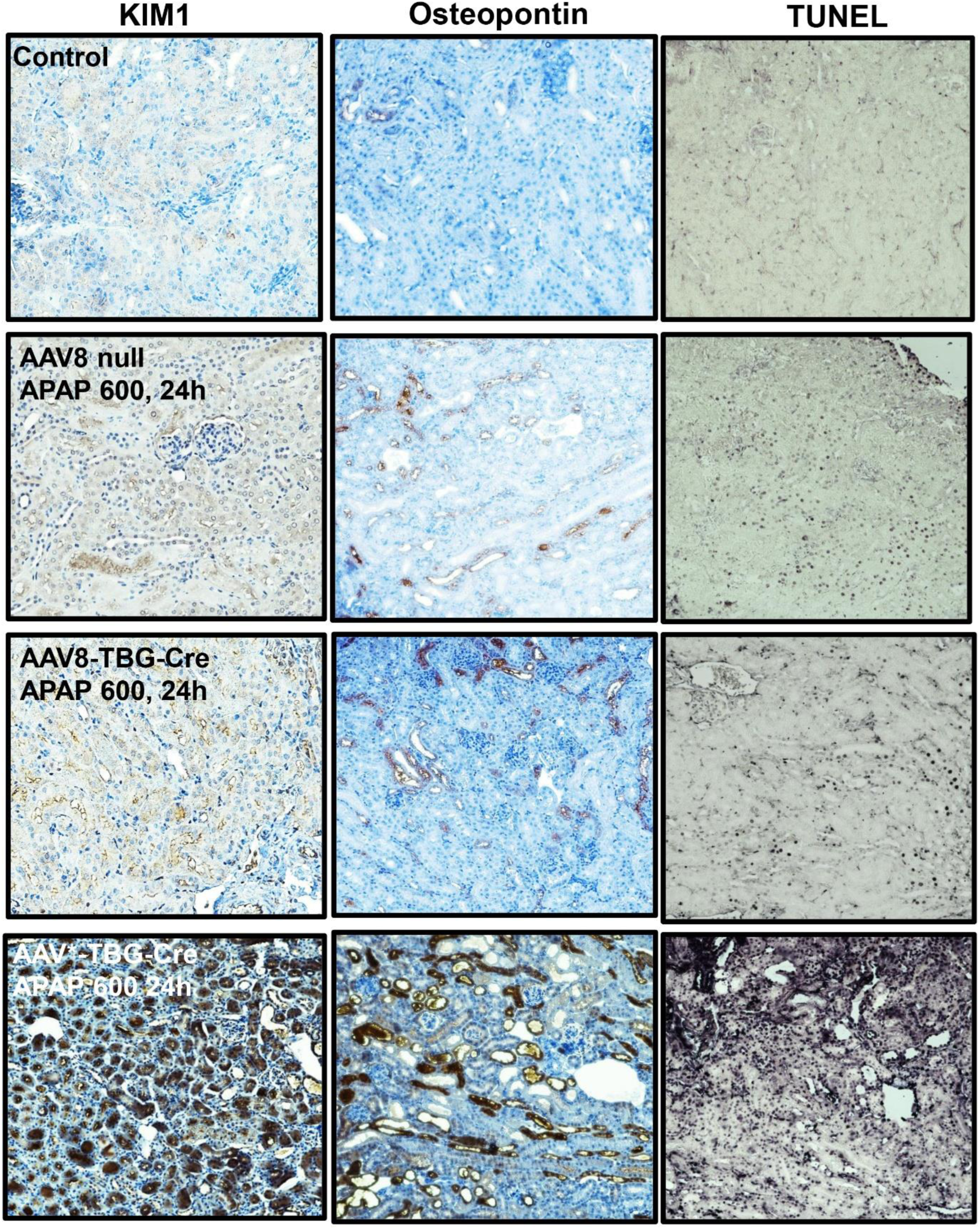
Temporal progression of APAP-induced renal injury following selective hepatic CYP2E1 deletion. Kidney sections were stained for KIM-1 (left column), osteopontin (middle column), and TUNEL (right column) to assess injury markers and cell death. Representative images are shown for control mice, AAV8-null mice treated with APAP (600 mg/kg, 24 h), AAV8-TBG-Cre mice treated with APAP 600mg/kg for 24 or 48 hours. Images are representative of n = 4–5 mice per group and captured at 20X magnification.

### 3.3. Absence of renal CYP2E1 protects female mice from APAP nephrotoxicity at 24 hours

APAP hepatotoxicity is driven by its bioactivation through cytochrome P450 enzymes, particularly CYP2E1, which generates the reactive metabolite NAPQI (McGill & Jaeschke, 2013). Prior studies reported that female mice are resistant to APAP-induced renal injury due to the lack of testosterone-dependent induction of renal CYP2E1 (Hoivik et al., 1995; Hu et al., 1993; Kennon-McGill & McGill, 2018; Pan et al., 1992). To validate this in our model, we examined CYP2E1 expression in kidney sections from male and female mice. Compared with males, females exhibited no significant increase in CYP2E1 mRNA levels after APAP overdose (Figure 5A, B) and no detectable CYP2E1 protein by immunohistochemistry (Figure 5C). This biological difference provides a model in which renal APAP bioactivation is inherently minimized, allowing us to investigate kidney-intrinsic injury pathways without the influence of CYP2E1-driven local NAPQI formation. To characterize organ-specific injury in females, mice were treated with 600 mg/kg APAP and evaluated at 24 hours. Serum ALT levels rose sharply to approximately 6,000 U/L (Figure 6A), indicating substantial hepatocellular damage. BUN levels increased to ∼40 mg/dL (Figure 6B), demonstrating concurrent early renal dysfunction. In contrast, creatinine level did not increase significantly (Figure 6C). PAS staining of the kidneys showed well-preserved proximal tubular morphology, with only rare TUNEL-positive nuclei detected in the cortex, indicating minimal early tubular injury in female mice after severe APAP overdose (Figure 6D).

**Figure 5:**
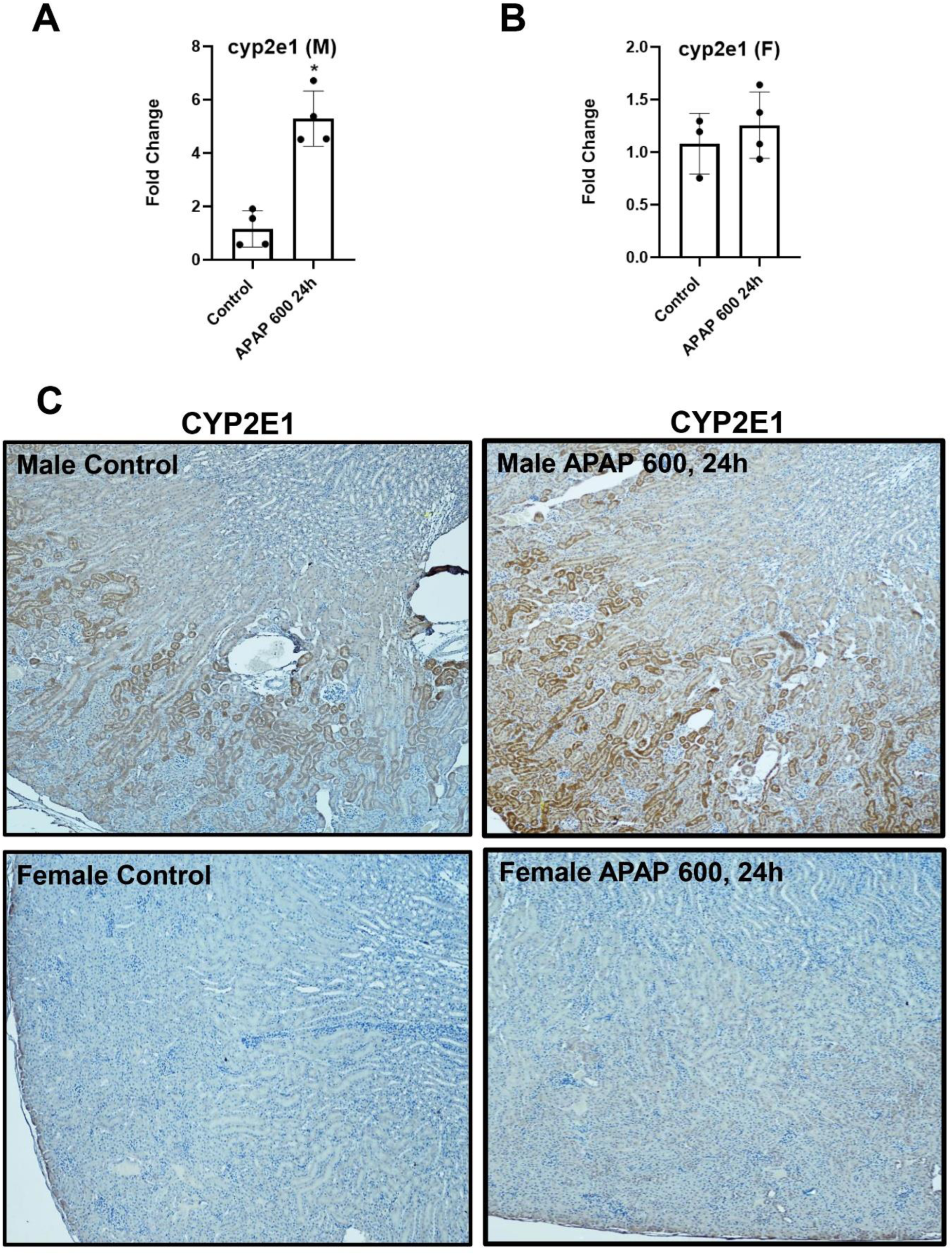
Sex-dependent renal CYP2E1 expression in response to APAP. Relative renal CYP2E1 mRNA expression in male (A) and female (B) C57BL/6J mice 24 hours after acetaminophen (APAP; 600 mg/kg) administration. (C) Representative kidney sections stained for CYP2E1 from male and female C57BL/6J mice under control conditions and 24 hours following acetaminophen (APAP; 600 mg/kg) administration. Images are representative of n = 4–5 mice per group and were captured at 10X. Bars represent means ± SEM for n = 3–4 mice per group. *p < 0.05 compared to control.

**Figure 6:**
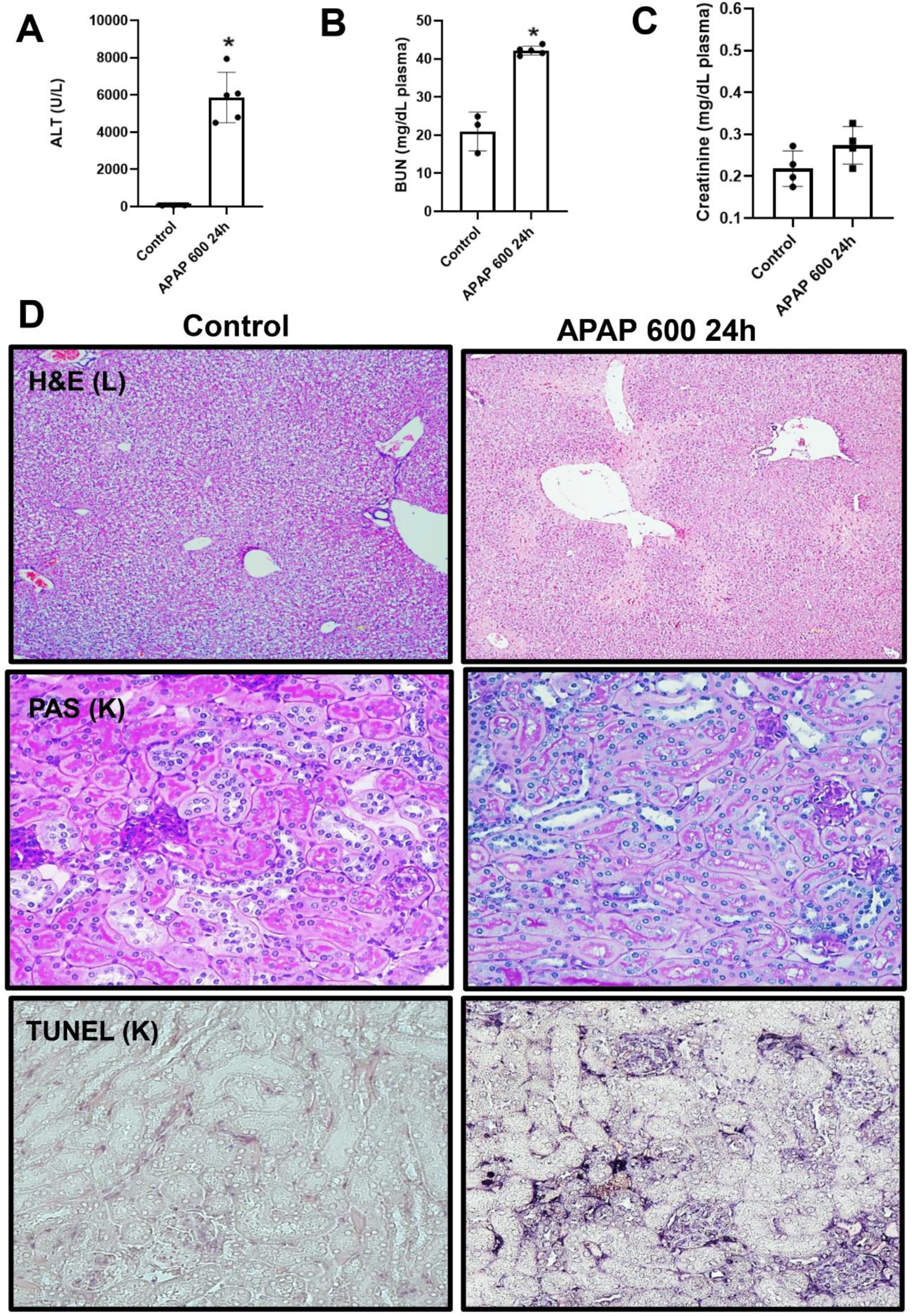
Lack of significant acute kidney injury in female mice following APAP 600 mg/kg administration. Female C57BL/6J mice received a single intraperitoneal injection of APAP (600 mg/kg) and were sacrificed 24 hours later. (A) Plasma ALT activities, (B) plasma BUN levels, and (C) plasma creatinine levels, (D) representative histological images including liver H&E staining (10X), kidney PAS staining (40X), and kidney TUNEL staining (40X). Bars represent means ± SEM for n = 4–5 mice per group. *p < 0.05 compared to control.

### 3.4. Female mice exhibit no detectable protein adducts after APAP treatment

Female mice were treated with 600 mg/kg APAP and analyzed at 3, 6, and 24 hours. Plasma ALT levels were markedly elevated at all time-points, confirming progressive hepatocellular injury (Figure 7A). Plasma BUN and creatinine levels increased as early as 3 h post-treatment. While BUN levels remained elevated at 24 h, creatinine levels returned to baseline by this time (Figure 7B,C). Given that hepatic APAP–protein adduct formation is known to be dose- and time-dependent and to occur rapidly after NAPQI generation (McGill et al., 2013), early kidney time points were included to determine whether transient adduct formation also occurs in the kidney. No APAP-protein adducts were detected in kidney homogenates at any time points (data not shown), indicating minimal renal bioactivation in females and suggesting that kidney injury does not arise from local CYP-dependent APAP metabolism. Consistent with these findings, immunohistochemical analysis of kidney injury markers revealed minimal induction in KIM1 or OPN expression in APAP-treated females at 24 hours compared with controls (Figure 7D).

**Figure 7:**
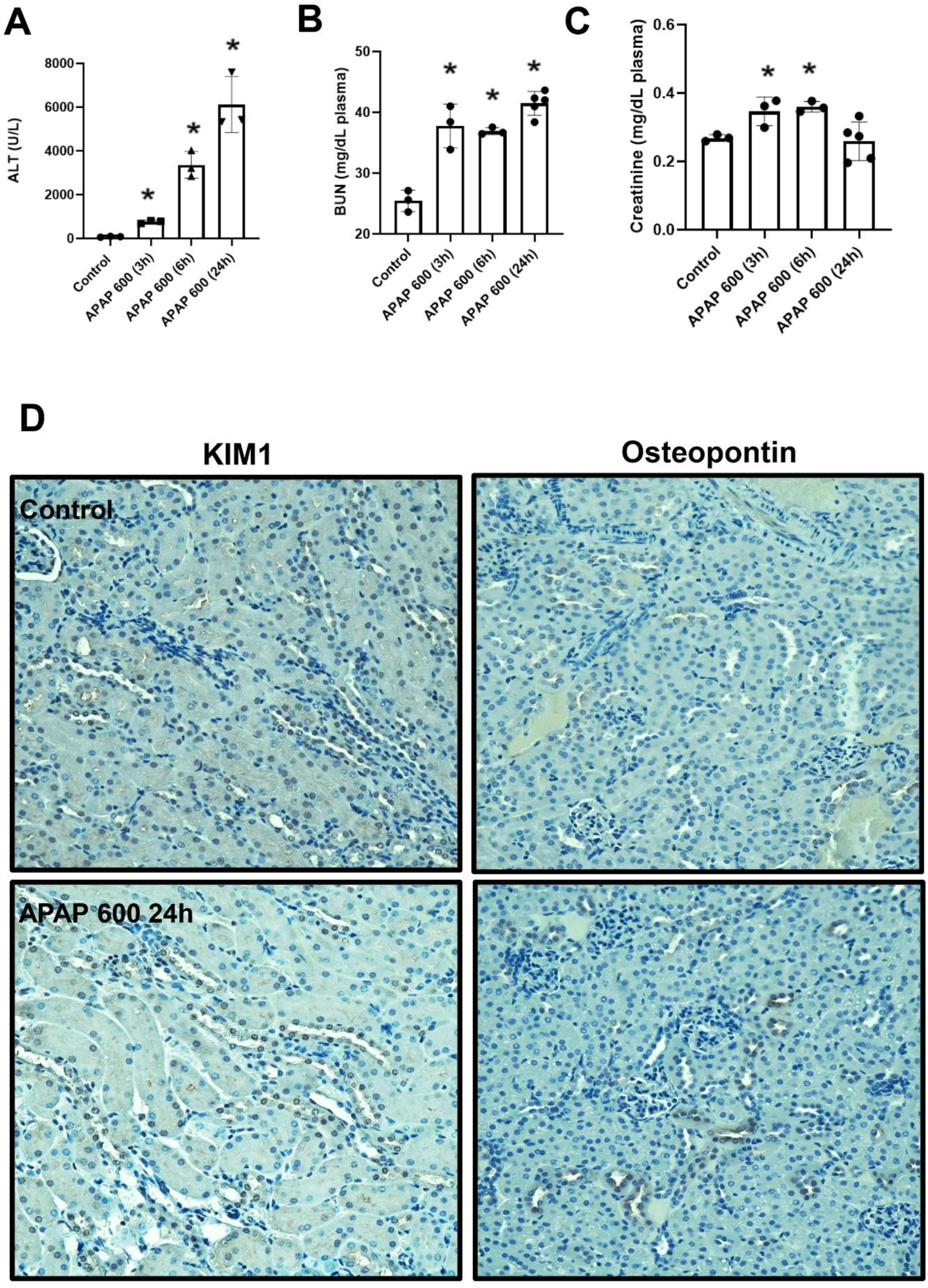
Liver and kidney injury biomarkers in female mice following APAP administration. Female C57BL/6J mice received a single intraperitoneal injection of APAP (600 mg/kg) and were sacrificed at 3, 6, or 24 hours. (A) Plasma ALT activities, (B) plasma BUN levels, (C) plasma creatinine levels at each time point, (D) representative kidney sections stained for KIM-1 and osteopontin from control and APAP-treated mice (600 mg/kg, 24 h). Images are representative of n = 4–5 mice per group and were captured at 40X magnification. Bars represent means ± SEM for n = 4–5 mice per group. *p < 0.05 compared to control.

### 3.5. CYP2E1-Independent renal injury progresses in female mice even after hepatic protection

As shown in earlier figures, female mice naturally lack renal CYP2E1, allowing APAP-induced kidney injury to be examined without local bioactivation. By additionally deleting hepatic CYP2E1 using AAV8-TBG-Cre in CYP2E1^flox/flox^ females, we generated a model in which both renal and hepatic CYP2E1 pathways are eliminated. This dual-deficiency system enables direct investigation of CYP2E1-independent mechanisms of kidney injury and, importantly, allows evaluation of renal injury beyond 24 hours without mortality from hepatic failure. Using this model, we next assessed liver and kidney injury markers in AAV8-TBG-Cre-treated female mice following APAP overdose. As expected, plasma ALT levels were markedly elevated in null mice at 24 hours (∼6000 U/L), indicating severe hepatocellular injury, whereas AAV8-TBG-Cre–treated mice showed >85% reduced ALT activity at both 24 and 48 hours (Figure 8A). In contrast, BUN levels rose to ∼40 mg/dL and creatinine also significantly increased across all groups (Figure 8B,C), indicating persistent renal dysfunction despite substantial protection from hepatic injury. CYP2E1 immunohistochemistry confirmed efficient and widespread depletion of hepatic CYP2E1 in AAV8-TBG-Cre-treated mice (Figure 8D). Consistent with this, AAV8-null-treated mice exhibited extensive hepatic necrosis at 24 hours, whereas AAV8-TBG-Cre treated mice showed moderate centrilobular necrosis at both 24- and 48-hours following APAP treatment. In contrast, kidney histology demonstrated progressive injury at 48 hours, evidenced by marked cytoplasmic vacuolization in proximal tubular cells (black arrows), while TUNEL staining showed abundant apoptotic nuclei, indicating ongoing tubular cell death (Figure 9). Further assessment of renal injury markers revealed strong KIM1 induction and increased osteopontin expression in the renal cortex at 48 hours (Figure 9), confirming acute kidney injury. Together, these findings verify effective hepatic CYP2E1 deletion and demonstrate that kidney injury continues to progress at 48 hours despite extensive protection from APAP-induced liver injury.

**Figure 8:**
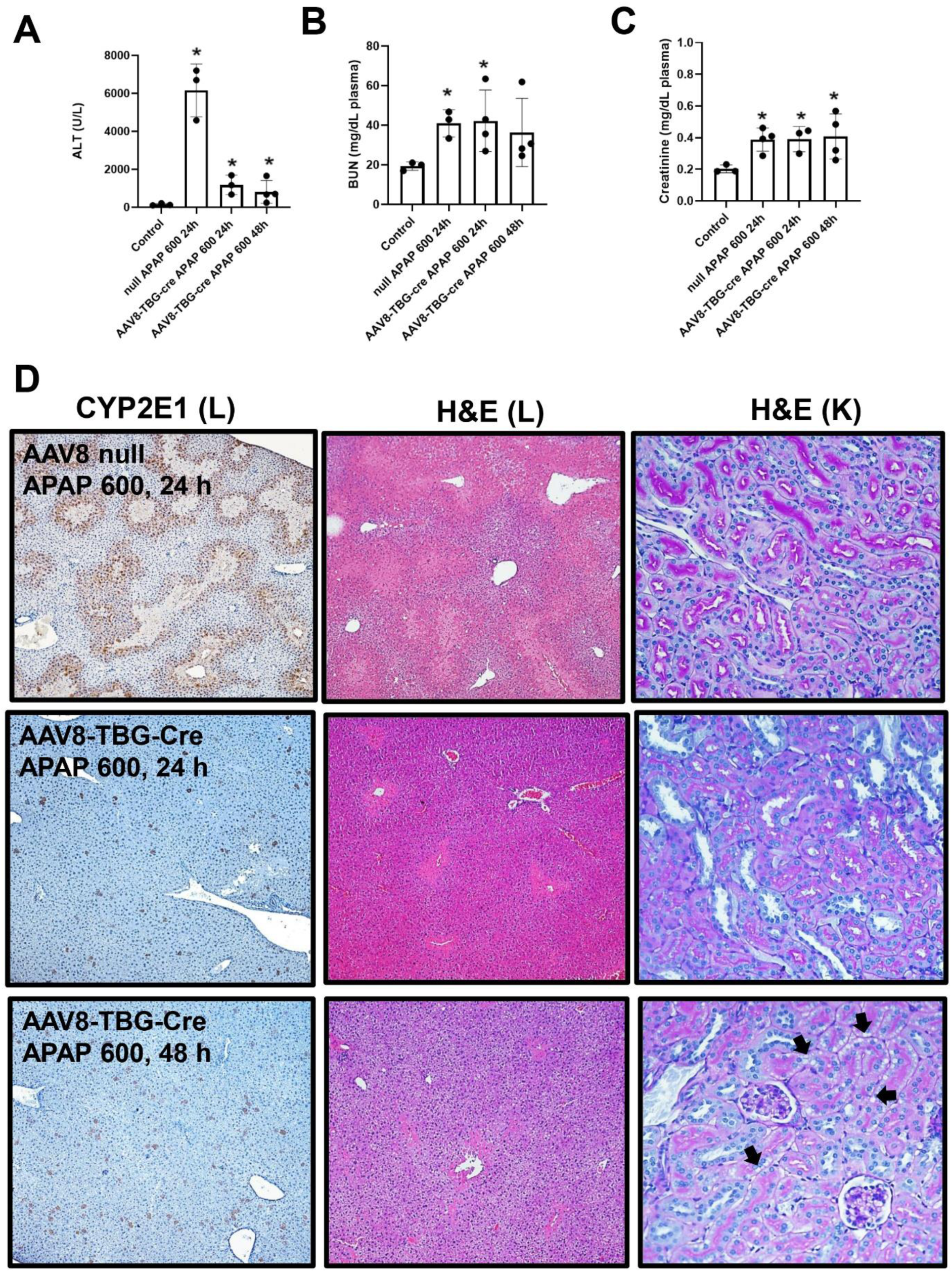
Delayed and progressive acute kidney injury in female mice lacking hepatic and renal CYP2E1. Female C57BL/6J mice received AAV8-null or AAV8-TBG-Cre vectors and, two weeks later, a single intraperitoneal injection of APAP (600 mg/kg). Mice were sacrificed 24 or 48 hours after APAP administration. (A) Plasma ALT activities, (B) plasma BUN levels, (C) plasma creatinine levels, (D) representative liver and kidney sections, including liver CYP2E1 immunohistochemistry (10X), liver H&E staining (10X), and kidney PAS staining (40X), from AAV8-null mice treated with APAP (600 mg/kg, 24 h), AAV8-TBG-Cre mice treated with APAP (600 mg/kg) for 24 or 48 hours. Black arrows indicate vacuolar changes observed at 48 hours. Images are representative of n = 4–5 mice per group. Bars represent means ± SEM for n = 4–5 mice per group. *p < 0.05 compared to control.

**Figure 9:**
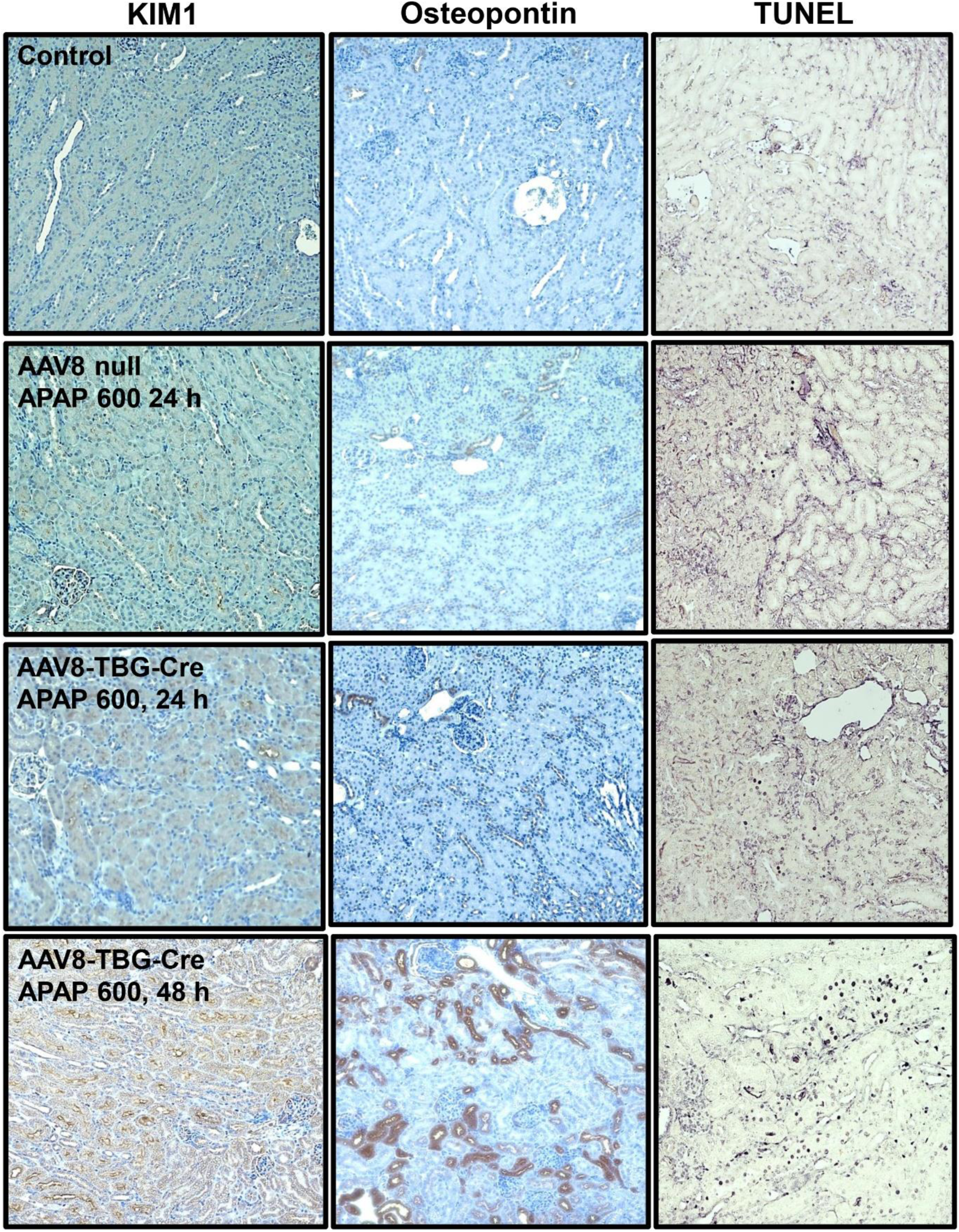
Temporal progression of APAP-induced renal injury in female mice. Kidney sections were collected at 24 or 48 hours after APAP administration and stained for KIM-1 (left column), osteopontin (middle column), and TUNEL (right column). Representative images are shown from control, AAV8-null mice treated with APAP (600 mg/kg, 24 h), and AAV8-TBG-Cre mice treated with APAP (600 mg/kg) for 24 or 48 hours. Images are representative of n = 4–5 mice per group and were captured at 20X magnification.

### 3.6. Evaluation of the role of APAP deacetylation in acute kidney injury

As demonstrated in previous experiments, deletion of CYP2E1 in both liver and kidney did not completely prevent renal injury, as substantial AKI still developed at 48 h following APAP administration. These findings indicate the involvement of additional CYP-independent mechanisms in APAP-induced nephrotoxicity. One potential pathway is the deacetylation of APAP to p-aminophenol, a known nephrotoxic metabolite associated with proximal tubular injury (Newton et al., 1985; Newton et al., 1982). To investigate the contribution of APAP deacetylation to kidney injury, AAV8-treated mice were pretreated with the carboxylesterase inhibitor TOTP (Emeigh Hart et al., 1991) 1 h before APAP administration (600 mg/kg) and sacrificed 48 h later. Plasma biomarkers and tissue-specific deacetylation activity were then assessed. TOTP pretreatment did not significantly alter plasma ALT activity (Figure 10A); in contrast renal injury markers were markedly attenuated at 48 hours, as evidenced by significant reductions in plasma BUN and creatinine levels (Figure 10B,C). These findings suggest a selective protective effect of TOTP on kidney injury. Consistent with these observations, ex vivo analysis of tissue homogenates demonstrated that inhibition of deacetylation with TOTP markedly reduced PAP formation in both kidney and liver (Fig. 10D,E), in agreement with previous reports (Emeigh Hart et al., 1991). Homogenates prepared from control mice generated high levels of PAP following incubation with 25 mM APAP for 1 h, as described previously (Emeigh Hart et al., 1991), whereas tissues from TOTP-treated mice exhibited minimal PAP formation, confirming effective inhibition of deacetylase activity. These findings confirm that TOTP effectively inhibits APAP deacetylation in both kidney and liver tissues. Histological evaluation further supported these findings. PAS staining revealed reduced tubular injury and cast formation in TOTP-pretreated mice compared with APAP-treated animals (Figure 10F). In addition, immunostaining for kidney injury markers showed decreased expression of KIM-1 and osteopontin in the TOTP + APAP group relative to the APAP alone-treated group (Figure 10F).

**Figure 10:**
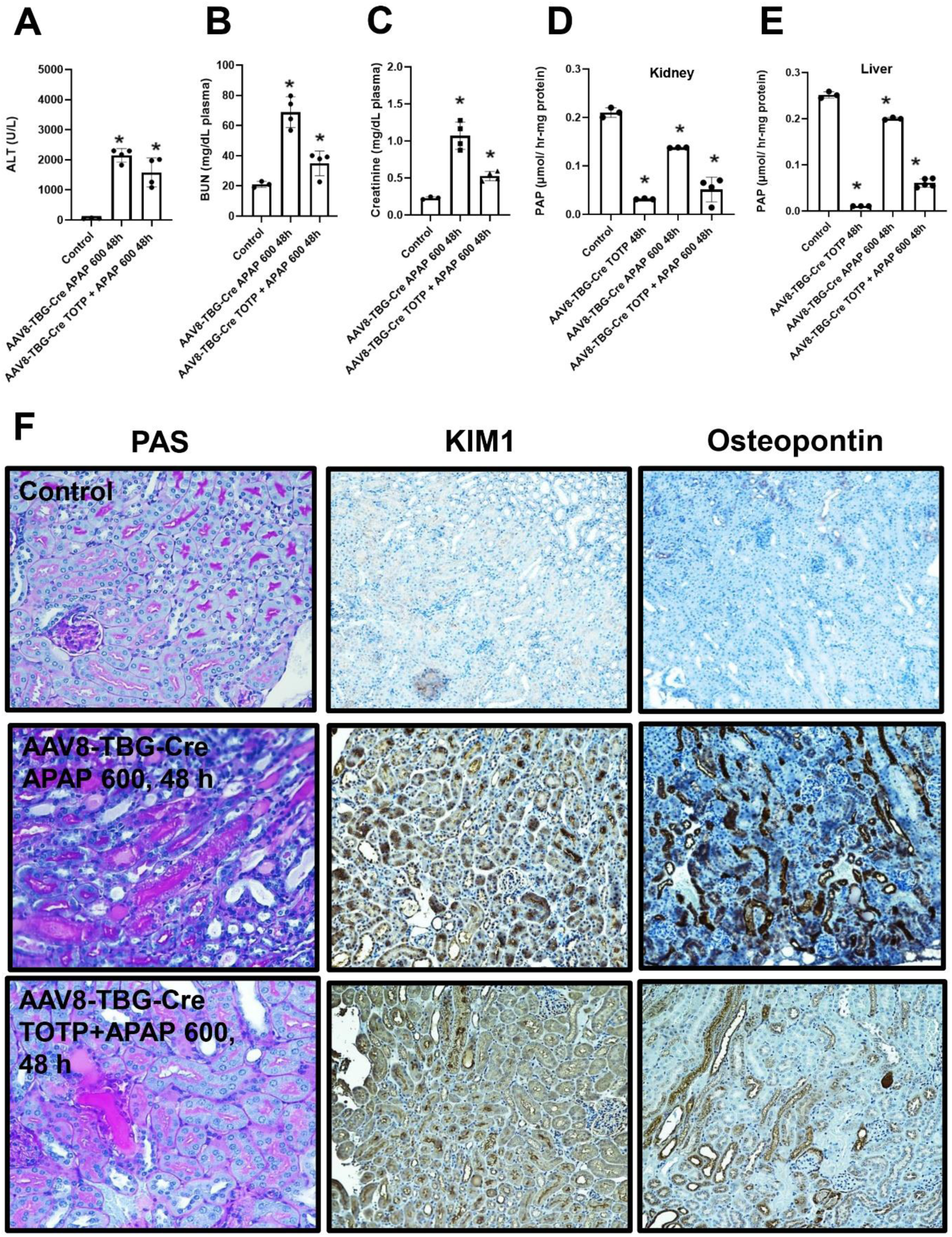
Effect of carboxylesterase inhibition on APAP-induced kidney injury and deacetylation. Male C57BL/6J mice were administered AAV8-TBG-Cre vectors and, after two weeks, received a single intraperitoneal injection of TOTP (125 mg/kg) of vehicle followed by APAP (600 mg/kg) 1 hour later. Mice were euthanized 48 hours after APAP treatment. Plasma biomarkers of injury were assessed, including (A) ALT activity, (B) BUN levels, and (C) creatinine levels. For deacetylation analysis, liver and kidney tissues were collected from vehicle-, TOTP-, APAP-treated, and TOTP + APAP-treated groups. Kidney (D) and liver (E) homogenate supernatants were incubated with 25 mM APAP for 1 hour at 37°C, and PAP formation was quantified. Representative kidney histology is shown by PAS staining (40×) for control, AAV8-TBG-Cre + APAP (48 h), and AAV8-TBG-Cre + TOTP + APAP (48 h) groups. Kidney injury markers were further evaluated by immunostaining for KIM-1 and osteopontin in sections collected at 48 hours post-APAP. Representative images are shown for control, AAV8-TBG-Cre + APAP (600 mg/kg, 48 h), and AAV8-TBG-Cre + TOTP + APAP (600 mg/kg, 48 h) groups. Images are representative of n = 3–4 mice per group and were acquired at 40X for PAS and 20X for KIM1 and osteopontin. Bars represent means ± SEM for n = 3–4 mice per group. *p < 0.05 compared to control.

## 4 Discussion

### 4.1 Kidney injury progresses independently of hepatic CYP2E1 bioactivation and continues to worsen over time

A central goal of this study was to determine whether APAP-induced kidney injury depends on the canonical hepatic bioactivation pathway mediated by CYP2E1. To address this question, we used AAV8-TBG-Cre to selectively delete hepatic CYP2E1 in male and female mice. This approach effectively reduced CYP2E1 expression and substantially attenuated liver injury, thereby creating a model in which renal outcomes could be evaluated in the absence of hepatic NAPQI formation. Our findings show that both male and female CYP2E1-floxed mice treated with AAV8-TBG-Cre and subsequently with APAP (600 mg/kg) developed evidence of kidney dysfunction and injury at 24 hours, despite being mainly protected from liver damage. By 48 hours, male mice progressed to marked renal dysfunction and proximal tubular damage. Female mice exhibited minimal renal injury at 24 hours but developed clear kidney injury by 48 hours. Together, these findings show that APAP overdose results in progressive, time-dependent kidney injury even after selective hepatic deletion of CYP2E1 by AAV8-TBG-Cre, demonstrating that renal injury can arise independently of hepatic APAP bioactivation. This conclusion is consistent with our previous study showing that manipulation of organ GSH content can affect renal and hepatic injury independent of each other (Etemadi et al., 2026).

These findings further align with prior studies indicating that renal APAP metabolism and toxicity occur via mechanisms distinct from the liver (Akakpo et al., 2023; Hart et al., 1994, 1995; Lorz et al., 2004; Mour et al., 2005). In the kidney, CYP2E1 is predominantly localized to the ER of proximal tubular cells. In contrast, in hepatocytes, CYP2E1 is found in both the ER and at higher levels within mitochondria (Akakpo et al., 2023), suggesting fundamental differences in subcellular bioactivation and downstream injury signaling. Importantly, kidney injury has also been shown to be exacerbated by APAP cysteine, a glutathione-derived metabolite, in the absence of concomitant liver injury, further supporting the concept that kidney-specific metabolic and non-canonical pathways can drive renal toxicity independently of hepatic NAPQI formation (Stern et al., 2005).

### 4.2 Renal injury in the absence of CYP2E1-Mediated APAP Bioactivation

Our previous work showed that renal APAP bioactivation in male mice depends on CYP2E1, and that pharmacological inhibition with fomepizole prevents both renal adduct formation and subsequent injury within 24 hours (Akakpo et al., 2020, 2023). In contrast, female mice showed no detectable APAP–protein adducts in the kidney at 3, 6, or 24 hours after APAP administration, which indicates that no other renal P450 isoform contributes to local NAPQI formation during this early phase. To further investigate renal mechanisms, female mice were treated with AAV8-TBG-Cre to delete hepatic CYP2E1. This intervention reduced liver injury, extended survival, and allowed evaluation of renal outcomes beyond 24 hours. Under these conditions, the female mice developed pronounced kidney injury by 48 hours. Collectively, these findings demonstrate that the delayed renal injury observed in AAV8-TBG-Cre–treated female mice can occur through mechanisms independent of CYP-mediated APAP bioactivation, highlighting the presence of alternative, non–P450-driven pathways contributing to late-stage kidney damage.

### 4.2.1 PAP formation as a CYP-independent mechanism of renal cell death

One potential CYP-independent mechanism contributing to APAP-induced acute kidney injury involves the deacetylation of APAP to PAP (Newton et al., 1982, 1985). While APAP hepatotoxicity is classically attributed to CYP-mediated formation of NAPQI, both hepatic and renal tissues possess deacetylase activity capable of generating PAP, providing an alternative metabolic pathway (Carpenter and Mudge, 1981; Miyakawa et al., 2015; Newton et al., 1985; Newton et al., 1982). Importantly, studies have demonstrated that APAP can induce P450-independent cytotoxicity at higher concentrations, with PAP formation contributing to delayed cell injury, supporting the concept that non-CYP pathways may become more prominent under conditions of high exposure (Miyakawa et al., 2015). Given that a severe APAP dose was used in our AKI model, this provided a strong rationale to further investigate the contribution of this pathway. In the kidney, Newton and colleagues established that PAP is a potent nephrotoxin and that inhibition of APAP deacetylation with Bis(p-nitrophenyl) phosphate (BNPP) reduces covalent binding and attenuates renal injury, implicating PAP as a mediator of APAP-induced nephrotoxicity (Newton et al., 1982, 1985).

However, the contribution of this pathway appears to be species- and strain-dependent. Studies in CD-1 mice have shown that inhibition of deacetylation did not significantly attenuate APAP-induced nephrotoxicity at the 12-hour time point, suggesting that PAP formation may not be a dominant mechanism in this model at earlier time points (Emeigh Hart et al., 1991). Because APAP-induced more severe acute kidney injury is a delayed event that typically develops beyond 24 hours, we investigated the potential contribution of PAP formation at later time points. In our study, inhibition of APAP deacetylation using the deacetylase inhibitor TOTP resulted in partial protection against kidney injury at 48 hours. This was evidenced by approximately 50% reductions in plasma BUN and creatinine levels, along with significant decreases in KIM-1 and osteopontin expression. In addition, histological analysis revealed marked attenuation of tubular dilation and cast formation, key features of severe renal injury. These findings indicate that inhibition of PAP formation can mitigate the progression of AKI during the delayed phase of injury, but inhibition of this pathway alone is insufficient to completely prevent injury.

### 4.2.2 Prostaglandin endoperoxide synthase (PGES) and N-deacetylase

Another potential CYP-independent mechanism of APAP-induced renal cell death involves non-P450 metabolism within the kidney. Studies have reported that in the renal medulla, PGES can oxidize APAP to reactive intermediates capable of inducing oxidative stress and tissue injury. Additionally, APAP can be deacetylated by renal N-deacetylase to yield PAP, a nephrotoxic metabolite which can bind to sulfhydryl groups and generate reactive oxygen species (Carpenter and Mudge, 1981; Duggin, 1996; Kennon-McGill and McGill, 2018; Sciskalska et al., 2015). These alternative metabolic routes may be particularly relevant in kidney regions (such as the medulla) with inherently low CYP450 activity (Mohandas et al., 1981) offering a plausible basis for delayed renal injury, especially in female mice, where classic CYP-mediated bioactivation is absent. More studies are required to evaluate injury mechanisms in detail.

## 5. Summary

This study established that APAP-induced kidney injury progresses through mechanisms that are independent of hepatic CYP2E1–mediated bioactivation and continues to worsen over time. By selectively reducing hepatic CYP2E1, we eliminated liver-derived NAPQI formation and significantly reduced hepatotoxicity, thereby enabling survival beyond 24 hours. As shown in our data, renal injury still emerged at 24 hours and progressed to substantial dysfunction and tubular damage by 48 hours in both sexes. The absence of renal APAP–protein adducts in female mice further confirms that delayed kidney injury can occur without local CYP-mediated activation, pointing instead to alternative, non–P450 pathways driving late-stage toxicity. Importantly, our findings provide evidence that PAP formation contributes to this delayed, CYP-independent injury. Inhibition of APAP deacetylation with the deacetylase inhibitor TOTP resulted in partial protection at 48 hours, reducing BUN and creatinine levels, tubular injury markers, and histological evidence of damage. However, incomplete protection indicates that PAP-mediated toxicity represents only one component of a multifactorial injury process. Together, these findings demonstrate that APAP-induced AKI is driven by kidney-intrinsic, CYP-independent mechanisms, including PAP formation, that become more prominent during the delayed phase of injury. These results highlight the importance of kidney-specific therapeutic strategies, as renal injury is a critical determinant of outcome in APAP-induced acute liver failure.

## Acknowledgement

This work was funded in part by the National Institute of Diabetes and Digestive and Kidney Diseases (NIDDK) grants R01 DK102142 and DK125465, and National Institute of General Medical Sciences (NIGMS) funded Liver Disease COBRE grants P20 GM103549 and P30 GM118247. Y.E. was supported in part by the Biomedical Research Training Program (BRTP) at the University of Kansas Medical Center. We thank Jephte Akakpo for technical support in performing HPLC analysis to detect protein adducts.

## Credit authorship contribution statement

**Yasaman Etemadi**: Writing – original draft, Methodology, Investigation, Data curation. **Timothy A. Fields**: Writing – review & editing, Investigation, Formal analysis. **Anup Ramachandran**: Writing – review & editing, Visualization, Validation, Investigation, Funding acquisition, Data curation. **Hartmut Jaeschke**: Writing – review & editing, Validation, Supervision, Project administration, Funding acquisition, Conceptualization.

## ABBREVIATIONS

AAV8: adenovirus serotype 8
AKI: acute kidney injury
ALF: acute liver failure
ALT: alanine aminotransferase
APAP: acetaminophen
BUN: blood urea nitrogen
Cyp2E1: cytochrome P450 2E1
GSH: glutathione
H&E: hematoxylin & eosin
KIM-1: kidney injury marker-1
NAC: N-acetyl-L-cysteine
NAPQI: N-acetyl-p-benzoquinone imine
OPN: osteopontin
PAP: p-aminophenol
PAS: Periodic Acid-Schiff stain
TBG: thyroid hormone-binding globulin
TOTP: tri-o-tolyl phosphate
TUNEL: terminal deoxynucleotidyl transferase-mediated dUTP nick end-labelling assay

